# Genetic Dissection of Haploid Male Fertility in Maize (*Zea mays L*.)

**DOI:** 10.1101/318386

**Authors:** Jiwei Yang, Haochuan Li, Yanzhi Qu, Qiong Chen, Jihua Tang, Thomas lübberstedt, Zonghua Liu

## Abstract

Haploid genome doubling is a key limiting step of haploid breeding in maize. Spontaneous restoration of haploid male fertility (HMF) provides a method by which costs can be saved and which does not require the use of toxic chemicals, in contrast to the artificial doubling process. To reveal the genetic basis of HMF, haploids were obtained from the offspring of 285 F_2:3_ families, derived from the cross Zheng58× K22. The F_2:3_ families were used as female donor and YHI-1 as the male inducer line. The rates of HMF from each family line were evaluated at two field sites over two planting seasons. Quantitative trait loci (QTL) for HMF were identified using a genetic linkage map containing 157 simple sequence repeat (SSR) markers. QTL for HMF displayed incomplete dominance. Transgressive segregation of haploids from F_2:3_ families was observed relative to haploids derived from the two parents of the mapping population. A total of nine QTL were detected, which were distributed on chromosomes 1, 3, 4, 7, and 8. Three QTL, *qHMF3b*, *qHMF7a*, and *qHMF7b* were detected in both locations, respectively. In our mapping population, HMF was controlled by three major QTL. These QTL could be useful to predict the ability of spontaneous haploid genome doubling in related breeding materials, and to accelerate the haploid breeding process by introgression or aggregation of those QTL.

## Introduction

Developing homozygous lines is a key step in maize breeding programs. In the traditional process, about six generations are needed to develop homozygous lines by continuous selfing, which is a time-consuming and expensive process [1]. The use of maize haploid plants provides a rapid and efficient method to develop homozygous lines [2]. Producing haploid plants in vivo has become a routine process and has been adopted widely for maize breeding during the past decade [3,4]. Doubled haploid (DH) technology has gradually become one of the three core technologies of modern breeding programs, along with transgenic and molecular marker-assisted breeding technology [5]. Moreover, DH technology enables opportunities for characterizing and utilizing the genetic diversity present in gene bank accessions of maize [6,7].

The DH process in maize includes three steps: production of haploids, haploid genome doubling, and DH line development and application. With the development of inducers such as WS14 from the cross W23 and Stock 6 [8], Zarodyshevy Mark Saratovsky, ZMS [9], China Agricultural University High Oil Inducer, CAUHOI [10], Moldovian Haploid Inducer [11], RWS from WS14 and KEMS [12], UH400 inducer of University Hohenheim [13], No. 3 inducer of Jilin Academy of Agricultural Sciences, JAAS3 [14] and No. 5 inducer of China Agricultural University CAU-5 [15], production of haploids has become increasingly efficient. In contrast, haploid genome doubling has become a limiting step of DH technology in large-scale applications.

Currently, artificial genome doubling using chemicals and spontaneous haploid genome doubling (SHGD) are mainly used for DH line production. During artificial genome doubling, chemicals are used, such as colchicine, trifluralin and pronamide, which are harmful to atmosphere, soil, and human health, due to their high toxicities [16]. In addition, artificial genome doubling is a complex process. Treated seedlings must be grown under controlled conditions, increasing breeding costs. Therefore, SHGD, where the fertility of maize haploids is restored under natural conditions without treatment, is a simpler and cheaper method. However, prerequisite is genetic variation for SHGD.

Both male and female floral organs have the capacity of SHGD [17]. Haploid female floral organs have a greater tendency for restoration of their fertility, with rates exceeding 90%. Chalyk [9] reported that 228 out of 234 ears of haploid plants (96%) carried kernels after pollination with pollen from diploid plants. Similarly, Liu and Song [18] found 93% of haploid ears to be naturally fertile. Therefore, the limiting factor for SHGD is, whether fertile pollen can be produced by haploid plants. Haploid male fertility (HMF) has been reported [13, 19–21]. HMF rates varied in different environments, and among genotypes, some genotypes with a zero HMF rate and other genotypes exceeding a rate of 10% [22]. Wu [23] reported no significant differences in HMF restoration rates between reciprocal crosses. Ren et al. [24] reported four QTL related to HMF and found a major QTL on chromosome 6. Their results suggested that HMF is affected by genetic background and environment.

So far, only few studies addressed the genetic basis of HMF [9,24,25]. Ren et al. [24] first reported the QTL related to HMF using two segregation populations from temperate germplasm Zheng58 crossed tropical Yu87-1 and Lancast germplasm 4F_1_. In this study, we used F_2:3_ families from the cross of inbred lines Zheng58 and K22, which both from temperate germplasm to determine HMF rates in two different environments over two years. The objectives of this study were to (i) characterize the mode of inheritance of HMF, and (ii) to detect QTL affecting HMF.

## Materials and Methods

### Plant Materials and Haploid Identification

Yu High Inducer No.1 (YHI-1), developed by Henan Agricultural University, was used as maternal inducer. A set of 285 F_2:3_ families from the cross between the two elite inbred lines Zheng58 and K22 from the same heterotic group germplasm, were used as female donors. Inbred line Zheng58 was developed by Henan Academy of Agricultural Science and has a low HMF rate (5.8%). In contrast, inbred line K22, developed by Northwest Agriculture and Forestry University, has a high HMF rate (56.4%). Induction crosses were produced in Hainan (N 18°21’, E109°10’; China) during the winter of 2013. YHI-1 is homozygous for the dominant marker gene *R-nj*. Purple coloration of embryo and endosperm was used as phenotypic marker to discriminate haploid and diploid kernels [26,27]. Putative haploid kernels with colorless embryos were planted in the field for verification based on plant vigor: haploid plants are short and weak, in contrast to vigorous hybrids. Thus, putative haploids with vigorous growth were eliminated as false positives.

### Field Treatment and Phenotypic Evaluation

Haploid plants from the F_2:3_ population and the two parents Zheng58 and K22 lines, were planted family-wise in fields at Zhengzhou experimental station, Henan Agricultural University (Zhengzhou, 113°42E, 34°480N) summer of 2014 and at Hainan experimental station (18°21N, 109°10E) winter of 2014, respectively. At each location, a completely randomized design was used. Experimental materials was planted in 4 m long rows with 0.6 m space between rows, at a density of 75,000 plants/ha. Standard agronomic practices such as irrigation, fertilization and weeding were used during each vegetation period, to ensure a uniform stand. During the pollen shedding and silking stages, plants with anthers exposed were classified based on the amount of pollen produced as male fertile haploids. The rate of HMF restoration was calculated by the formula as below:

HMF = (HMFN/N) × 100%;

Where, HMF is the rate of haploid male fertility; HMFN is the number of the haploid male fertile plants in each plot; these haploid plants with exposed anthers were able to produce viable pollen; N is the number of the total haploid plants in each plot.

Data analysis was performed in the SAS 8.2 statistical software package, using the PROC MIXED procedure [28]. The statistical model was as follows:

*Y_ij_*=*μ*+ *G_i_* + *L_j_* +*ε_ij_*

*Y_ij_* is the value of *i^th^* genotype at the *j^th^* location, *μ* is the overall population mean, *G_i_* is the effect of genotype, *L_j_* the effect of location, and *ε_ij_* the error term. All of the factors were treated as random effects.

### Genetic Map Construction and QTL Mapping

Leaf samples of the F_2_ population were collected at the seedling stage in the field, and the Sodium Laureth Sulfate (SLS) method [29] was used for DNA extraction. Simple sequence repeat (SSR) analysis was conducted as reported by Senior and Heun [30]. Polymorphisms between the two parent lines, Zheng58 and K22, were screened using 1200 pairs of SSR markers, distributed across the whole maize genome (http://www.maizegdb.org), and 157 SSR markers with distinct polymorphisms between the two parents were chosen. Linkage analysis was performed using MAPMAKER/EXP 3.0 [31,32]. QTL were detected using Win QTL Cartographer V2.5 software [33], based on composite interval mapping (CIM) fitting parameters for a targeted QTL in one interval, with a stepwise forward-backward regression analysis (Model 6 from Win QTL Cartographer V2.5). The genome was scanned in 2 cM intervals using regression analysis. Default values of 5 for the control markers and 10 for the window size were used. The threshold for the logarithm of odds (LOD) scores was estimated using permutation tests [34] with 1000 replications at a *P*=0.05 level of significance for an experiment wise Type I error.

The QTL notation followed the rules suggested by McCouch et al.[35], each QTL name was started with a lowercase ‘*q*’, then the trait name in capital letters, followed by a figure showing the chromosome number where the QTL was detected. If there were more than one QTL for the same trait on the same chromosome, a lowercase letter was added after the chromosome number to distinguish these QTL.

## Results

### Phenotypic Data Analysis of HMF

There were differences in HMF between the two parents in Zhengzhou and Hainan (Table 1). In Zhengzhou, the inbred line Zheng58 had a mean HMF rate of 5.3%, showing low fertility restoration, while the rate of line K22 was 57.1%. The level of HMF in Hainan is similar to Zhengzhou, the HMF rates of Zheng58 and K22 were 6.3% and 55.6% in Zhengzhou, respectively. The average rate of HMF for Zheng58 is 5.8%, while that of K22 is 56.4%. The mean HMF rate of the F_2:3_ population across both environments was slightly lower than the mid-parent value, but there is no significant difference between parent and population means. HMF presents a proximate continuous distribution in each location (S1 Fig), consistent with a normal distribution. The coefficient of Skewness is a measure for the degree of symmetry and the coefficient of Kurtosis is a measure for the degree of tailedness in the variable distribution [36,37]. Skewness and kurtosis coefficients in this study, respectively (*P*=0.56>0.05), were consistent with a normal distribution.

**Table 1.** Phenotypic analysis of fertility restoration rates in the haploid male plant parts of parents and their offspring populations in maize

| Location | Zheng58 | K22 | F1 | F<sub>2:3</sub> family lines |  |  |  |  |  |
| --- | --- | --- | --- | --- | --- | --- | --- | --- | --- |
|  |  |  |  | Minimum value(%) | Maximum value(%) | Mean | CV | Skewness | Kurtosis |
| Zhengzhou | 5.26 | 57.14 | 35.29 | 0 | 100 | 27.02 | 0.73 | 0.19 | 0.76 |
| Hainan | 6.25 | 55.56 | 36.36 | 0 | 100 | 30.93 | 0.78 | -0.41 | 0.51 |

HMF rate of the F_1_ (35.8%) between both parents exceeded the mean of both parents of 31.1% across both environments. Some of the F_2:3_ families transgressed the parents for HMF, the lowest and highest HMFR in population reached 0 and 90% (S1 and S2 tables). HMF rates differed significantly among genotypes and locations for the F_2:3_ population (Table 2).

**Table 2.** Variance analysis of haploid male fertility for the F_2:3_ populations in Hainan and Zhengzhou

| Sources | SS | df | MS | F value | F<sub>0.05</sub> | F<sub>0.01</sub> |
| --- | --- | --- | --- | --- | --- | --- |
| Family lines | 212242.23 | 284 | 747.33 | 3.41** | 1.22 | 1.32 |
| Locations | 2177.23 | 1 | 2177.23 | 9.93** | 3.87 | 6.72 |
| Error | 62280.21 | 284 | 219.3 |  |  |  |
| Total | 276699.67 | 569 |  |  |  |  |

### Molecular Marker Linkage Map

The molecular linkage map includes 157 markers for genotyping of the 285 F_2_ individuals (Fig 1). The linkage groups had a total length of 1927.1 cM and there was a mean distance of 12.3 cM between adjacent markers. The order of marker loci in the linkage map agreed well with that of the SSR bin map of the inter-mated B73×Mo17 population based on the AGI’s B73 RefGen_v2 sequence, (http://www.maizegdb.org), except for umc1841 (assigned to bin 7.03, but placed on chromosome 2 in our linkage map).

**Fig 1.**
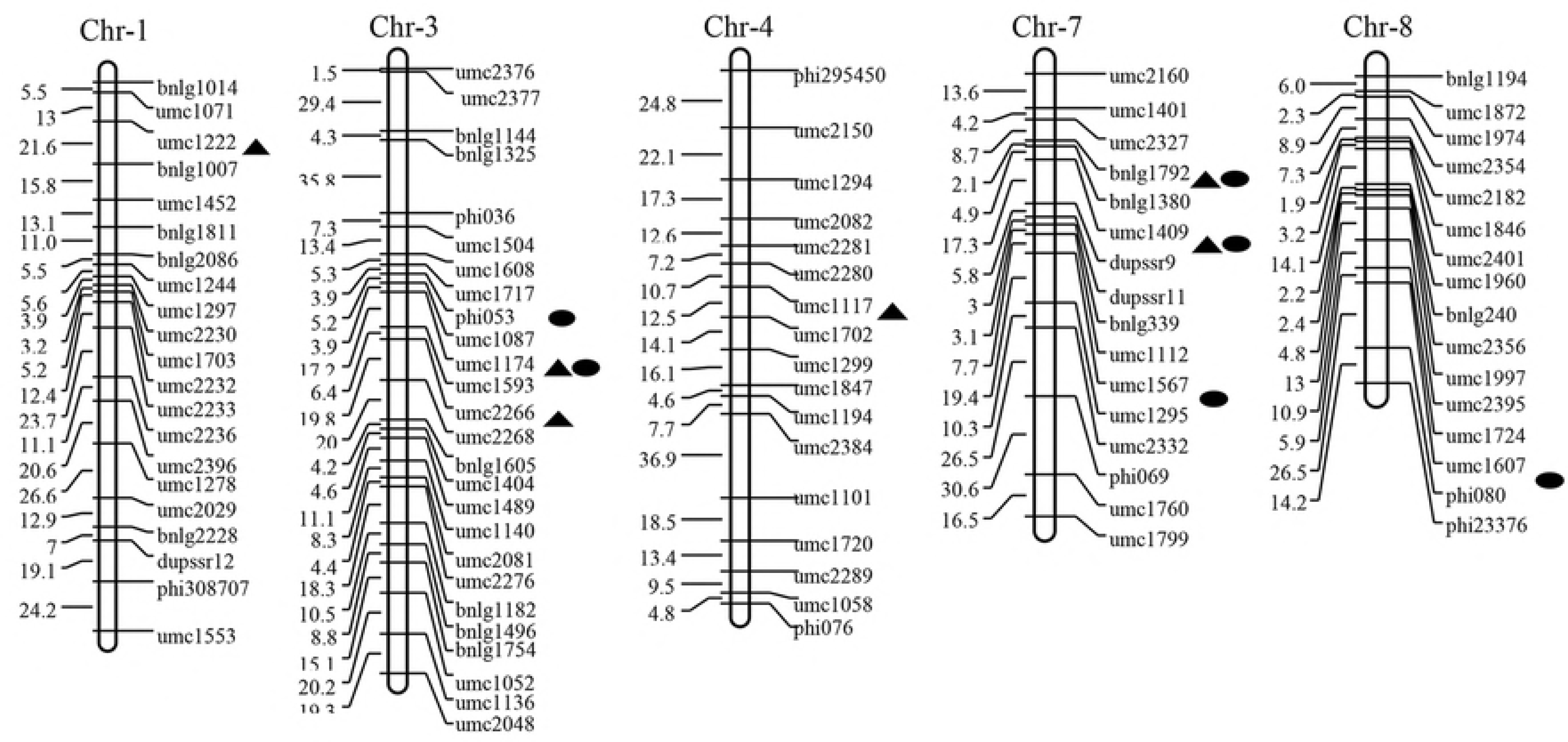
Chromosomal location of the QTLs used to assess haploid restoration of male fertility. Triangles denote an unconventional QTL detected in plants grown at the Hainan field site; ellipses denote a conventional QTL detected in plants grown at the Zhengzhou field site.

### QTL Analyses

Using CIM for QTL mapping analysis within and across both environments, 12 QTL were detected (Table 3). Six QTL, including *qHMF3a*, *qHMF3b*, *qHMF7a*, *qHMF7b*, *qHMF7c,* and *qHMF8*, were detected for Zhengzhou. The phenotypic contributions of individual QTL ranged from 6.3% to 12.2%, with a total contribution of 58.7%. For all six QTL, the favourable alleles came from inbred K22.

**Table 3.** Putative QTL detected for restoration of haploid male fertility for the F2:3 populations

| Location | QTL | Ranking-markers | Bin-locus<sup>a</sup> | Position<sup>b</sup> | LOD | A<sup>c</sup> | R<sup>2</sup> (%)<sup>d</sup> |
| --- | --- | --- | --- | --- | --- | --- | --- |
| Zhengzhou | *qHMF3a* | Phi053-umc1087 | 3.05 | 104.91 | 6.71 | -9.43 | 10.23 |
|  | *qHMF3b* | umc1174-umc1593 | 3.05 | 114.01 | 6.3 | -10.26 | 12.19 |
|  | *qHMF7a* | bnlg1792-bnlg1380 | 7.02 | 28.51 | 6.94 | -7.12 | 6.34 |
|  | *qHMF7b* | umc1409- dupssr9 | 7.01-7.02 | 41.51 | 7.17 | -8.93 | 10.24 |
|  | *qHMF7c* | umc1567-umc1295 | 7.03-7.04 | 74.41 | 6.6 | -9.36 | 11.19 |
|  | *qHMF8* | umc1607-phi080 | 8.07-8.08 | 98.91 | 5.12 | -7.98 | 8.54 |
| Hainan | *qHMF1* | umc1222-bnlg1007 | 1.02-1.02 | 32.51 | 2.64 | 6.63 | 3.23 |
|  | *qHMF3b* | umc1174-umc1593 | 3.05 | 120.01 | 7.79 | -6.52 | 3.32 |
|  | *qHMF3c* | umc2266-umc2268 | 3.06 | 141.61 | 8.94 | -7.98 | 5.17 |
|  | *qHMF4* | umc1117-umc1702 | 4.04-4.05 | 100.71 | 8.1 | -9.58 | 7.88 |
|  | *qHMF7a* | bnlg1792-bnlg1380 | 7.02 | 26.51 | 7.98 | -8.65 | 6.62 |
|  | *qHMF7b* | umc1409- dupssr9 | 7.01-7.02 | 43.51 | 7.42 | -9.56 | 8.13 |

<sup>a</sup>Bin locations of the flanking markers from the Maize GDB (http://www.maizegdb.org).
<sup>b</sup>Genetic map position, by cM.
<sup>c</sup> Additive effects estimated using QTL Cartographer.
<sup>d</sup>R<sup>2</sup> percentage of the phenotypic variance explained by the QTL.

At Hainan, six QTL for HMF were detected, including *qHMF1*, *qHMF3b*, *qHMF3c*, *qHMF4*, *qHMF7a*, and *qHMF7b*. The phenotypic contributions of individual QTL ranged from 3.2% to 8.1%, with a total contribution to phenotypic variance of 34.4%. The favourable alleles controlling HMF originated from inbred K22, except for *qHMF1* from Zheng58.

Three common QTL, *qHMF3b*, *qHMF7a*, *qHMF7b*, located between umc1174-umc1593 (chromosome 3), bnlg1792-bnlg1380 (chromosome 7), and umc1409- dupssr9 (chromosome 7), were detected across both locations. Their phenotypic contributions were 12.19%, 6.34% and 10.24%, at Zhengzhou, and 3.32%, 6.62% and 8.13%, respectively, at the Hainan site; again a slightly lower contribution rate (by 10.7%) of the three common QTL was observed in the plants from Hainan. All three of the common QTL were synergistic and were from the paternal inbred line K22. The results inferred that the actions of related QTL or genes varied with environment, at least to some degree.

## Discussion

There is no uniform standard to measure the characteristics of haploid fertility restoration. Kleiber et al. [17] and Ren et al. [24] scored anther emergence and classified haploids into a five-point scale based on (score 1) less than 5%, 6-20% (score 2), 21-50% (score 3), 51-75% (score 4), and 76-100% (score 5) anthers emerged on the tassel. Chalyk [9] and Geiger et al. [20] assessed shedding efficiency in their studies. In other studies, haploid inbred seed set has been used to determine male fertility [16]. To accurately assess haploid fertility restoration, the capacity of tassels to restore fertility (producing fertile pollen) and ear fertility restoration (bearing seed) should both be included [19]. Some anthers exposed in the tassel cannot produce viable pollen. Therefore, we combined both anther exposure and visual viable pollen to assess HMF.

Restoration of HMF is the main limiting factor for restoring haploid fertility, because haploid ears have shown high fertility rates of more than 90% [9,18,38]. Spontaneous restoration of HMF differed, when different sowing dates were used [39–41], and environment also affected HMF [42–44]. This may be related to temperature regimes or photoperiod, which influence gene expression. Liu and Song [18] reported that a negative (or positive) correlation tendency was shown between spontaneous restoration of HMF and temperature (or temperature difference between day and night) during early growth period of haploid plants. In a previous study, the rate of HMF in Hainan was higher than that in Zhengzhou, where the temperature difference between day and night is smaller than in Hainan.

In this study, the spontaneous restoration rates of HMF from the F_1_ and F_2:3_ generations were intermediate between the high parent line K22 and low parent line Zheng58. This suggests partial dominant inheritance of HMF. HMF of the different families from F_2:3_ population differed significantly according to variance analysis, the range of HMF was from 0 to 90% across both locations. Thus, both parent lines likely contain genes controlling HMF restoration. This was supported by QTL results. One QTL (*qHMF1*) originated from low parent line Zheng58 (negative additive effect), the other QTL from high parent line K22 (positive additive effect). It indicates that both parents perform differently for HMF depending on genetic backgrounds. However, the favorable HMF QTL can be aggregated in single lines to increase HMF. Consequently, the rate of HMF from some of families was higher than both parents and showed transgression, while for some other families had lower HMF than the parents(as low as 0%), because of negative locus aggregation. Therefore, it is possible to aggregate the positive alleles to enhance natural restoration ability of HMF.

There have been various mapping studies for haploid induction in maize [45–50], but only few investigated spontaneous haploid genome doubling. Wu et al. [16] reported a particular type of doubled haploids, named “early doubled haploids”, which were directly generated by in vivo haploid induction. It is likely that spontaneous doubling in embryo haploid (EH) only occurred during haploid embryo development after induction. However, early doubled haploids occurred at a frequency of 1-3.5%, which does not meet the demand for DH breeding at a large scale. Thus, HMF for haploid plants became of increasing interest. In a previous study, Wu [23] used 186 F_2:3_ families derived from a cross between Zheng58 (Reid heterotic group) and Chang7-2 (Tangsipingtou heterotic group) as female and CAU5 as male to obtain haploids from each family. Based on anther emergence score of haploids per se, eight QTL were detected on chromosomes 2, 3, 8, and 9. Only the locus on chromosome 8 was detected in both years. Ren et al. [24] reported four HMF QTL, *qhmf1*, *qhmf2*, *qhmf3*, and *qhmf4*, identified by segregation distortion. QTL detection was done in the selected haploid population derived from ‘Yu87-1/Zheng58’, and 48 recombinants were used to narrow the *qhmf4* locus down to an ~800 kb interval flanked by markers IND166 and IND1668. In this study, nine QTL for HMF were detected on chromosomes 1, 3, 4, 7, and 8, of which three QTL were detected in both field sites, even though they were grown during different seasons (winter and summer 2014). By comparison, the phenotypic contributions of *qHMF1*, and *HMF3b* (in Hainan) were lower than 5%, while the others contributed more than 5%. The detected HMF QTL in our study did not completely match those QTL reported previously (S3 Table). Based on physical coordinates from reference sequences, there is an overlap of HMF QTL with flanking markers umc2266-umc2268 (present study), bnlg1035-umc1528 [24], and umc1539-umc1528 [23], on chromosome3, as well as umc1997-dupssr14 [23] and umc1607-phi080 (present study) on chromosome 8. Up to now, 21 QTL related to HMF have been detected in the present and previous studies. A QTL on chromosome 3 was detected seven times, followed by QTL on chromosomes 2 and 7 detected three times each. These results imply that genes controlling HMF are distributed widely in germplasm of different genetic backgrounds.

Haploid male fertility was confirmed in this study as a quantitative trait controlled by many genes. QTL with major effect and stable expression are most important for MAS [51], we found three common loci for HMF by QTL mapping in both environments. These three QTL could be useful to enhance the spontaneous restoration ability of HMF by MAS to select individuals with favorable alleles, which can reduce the efforts for phenotypic selection. Furthermore, these molecular markers can be used to predict the ability of HMF in various breeding materials such as inbred lines, F_1_, F_2_, BC_1_ (backcross generation), etc. For the materials with high ability of HMF, doubled haploids (DH) lines will be produced by SHGD, while for those with poor ability of HMF, artificial genome doubling methods will be used to obtained more DH lines. In conclusion, novel QTL for HMF were detected in our study, which provides a base of understanding the genetics of HMF, and could be useful in guiding haploid doubling to increase the efficiency of haploid breeding programs and to accelerate maize breeding processes.

## Conclusions

Doubled haploid technology is the core factor that is limiting an increase in the speed, systematization and efficiency of the engineering processes employed during haploid breeding in maize. A more complete doubled haploid technology, based on production experiments is needed. Most of the methods require the use of chemical agents and the appropriate environment. This experiment shows that the spontaneous restoration ability of haploid male fertility (HMF) as a quantitative trait exists widely in maize germplasm with different genetic backgrounds, controlled by nuclear inherited and micro-effect polygenes and appeared incomplete dominance hereditary character. The male fertility restoration genes of the F_2:3_ population haploids from Zheng58 and K22 lines were studied using QTL mapping; three common loci were detected in plants grown at two locations, during different seasons. The results will allow a great improvement in the efficiency of promoting natural haploid doubling. It will provide some theoretical basis and practical experience about SGHD for haploid breeding technologies.

## Supporting information

**S1 Fig. Normal Q-Q plot for the rate of HMF from the F_2:3_ population in Zhengzhou and Hainan.**

**S1 Table. F_2:3_ families with higher HMF rate than K22(high parent)**

**S2 Table. F_2:3_ families with lower HMF rate than Z58 (low parent)**

**S3 Table. The detection of QTL for HMF in present and previous researches**

## Author Contributions

HL and ZL conceived and designed the research, JY, YQ and QC performed the experiments, JY analyzed the data, JT contributed to reagents and analysis tools, JY and HL wrote the manuscript, TL contributed to preparation of the manuscript. All authors read and approved the manuscript.

